# FANCD2 tunes the UPR preventing mitochondrial stress-­induced common fragile site instability

**DOI:** 10.1101/808915

**Authors:** Philippe Fernandes, Benoit Miotto, Claude Saint-Ruf, Viola Nähse, Silvia Ravera, Enrico Cappelli, Valeria Naim

## Abstract

Common fragile sites (CFSs) are genomic regions frequently involved in cancer-associated rearrangements. Most CFSs lie within large genes, and their instability relies on transcription- and replication-dependent mechanisms. Here, we uncover a role for the UBL5-dependent branch of the unfolded protein response pathway (UPR) in the maintenance of CFS stability. We show that genetic or pharmacological UPR activation induces CFS gene expression and concomitant relocalization of FANCD2, a master regulator of CFS stability, to CFSs. Furthermore, a genomic analysis of FANCD2 binding sites identified an enrichment for mitochondrial UPR transcriptional response elements in FANCD2 bound regions. We demonstrated that depletion of FANCD2 increases CFS gene transcription and their instability while also inducing mitochondrial dysfunction and triggering the activation of the UPR pathway. Depletion of UBL5, a mediator of the UPR, but not ATF4, reduces CFS gene expression and breakage in FANCD2-depleted cells. We thus demonstrate that FANCD2 recruitment and function at CFSs depends on transcription and UPR signaling, and in absence of transcription or UBL5, FANCD2 is dispensable for CFS stability. We propose that FANCD2 coordinates nuclear and mitochondrial activities by tuning the UPR to prevent genome instability.

## Introduction

Common fragile sites (CFSs) are genomic regions that are prone to form breaks and gaps within metaphase chromosomes after replication stress and drive genomic instability in the earliest steps of tumor development ^1^. CFS instability is cell-type dependent and relies on the cell replication and transcription programs ^2–4^. Notably, the transcription of very large genes encompassing CFSs can conflict with replication ^2^ and modify their replication dynamics ^5^, leading to their incomplete replication when cells enter mitosis. Incomplete replication of CFSs leads to the persistence of late replication intermediates that are processed by structure-specific endonucleases, inducing mitotic defects and genomic instability if not properly resolved in a timely manner ^6–8^. Despite their intrinsic instability, CFSs and their associated genes are conserved throughout evolution, hinting to a function of CFSs as sentinels of cellular stress ^9, 10^.

Among the many proteins involved in DNA replication/repair, the members of the FANC pathway (encoded by the *FANC* genes) are master regulators of CFS maintenance ^11^, with the FANC pathway being dysfunctional in individuals with Fanconi anemia (FA). FA is a rare chromosome instability disorder characterized by bone marrow failure, predisposition to acute myeloid leukemia and epithelial cancer, and hypersensitivity to DNA interstrand crosslinks (ICLs) and endogenous aldehydes^12 13, 14^. Chromosomal aberrations in FA patients are not random but occur preferentially at CFSs ^15–17^. FANCD2, the key activated target of the pathway, has been shown to relocalize at large genes encompassing CFSs after replication stress ^18, 19^ and form foci at CFSs during mitosis, where it cooperates with the helicase BLM (mutated in Bloom’s syndrome) to prevent chromosomal abnormalities ^20, 21^. In vivo, CFS instability can occur following physiological replication stress and is associated with impaired karyokinesis and megakaryocyte differentiation in *Fanca* −/− mice ^22^. Recent reports have highlighted the role of the FANC pathway in coordinating replication and transcription by preventing or resolving R-loops ^23, 24^. FANCD2 has been consistently shown to promote CFS replication by limiting R-loop formation ^25^. Therefore, failure to prevent or physiologically resolve R-loops and transcription-associated DNA damage may be the cause of the genomic instability that underlies the cancer predisposition of FA patients.

In addition to their nuclear functions, FANC proteins have been shown to play non-canonical roles in the regulation of mitochondrial function and redox metabolism ^26^. FANCD2 regulates mitochondrial energy metabolism by interacting with ATP5a ^27^, and Fancd2 has been shown to interact with components of mitochondrial nucleoid and to regulate mitochondrial gene transcription and translation in vivo ^28 29^. In addition, FANC proteins have been reported to regulate mitophagy by interacting with PARK2 ^30^, the product of the *PRKN* gene (also known as *Parkin*) encompassing the CFS FRA6E, which is mutated in Parkinson disease and is involved in mitochondrial quality control ^31^.

Mitochondrial dysfunction is an important effector of the FA cellular and clinical phenotype ^32, 33^, which is underscored by the fact that the tumor incidence and the hematopoietic defects in *Fanc*-deficient mice can be improved by antioxidant treatments ^34, 35^. However, whether these two independent functions in mitochondrial homeostasis and genome stability are mechanistically linked has remained unknown.

Mitochondria are key organelles that regulate many aspects of cellular metabolism, including energy production and nucleotide and amino acid metabolism. They are bounded by a double membrane system with 4 distinct functional compartments, the outer and inner membranes, the intermembrane space and the matrix, and maintenance of the protein-folding environment in each compartment is fundamental for proper organelle function ^36^. The components of the respiratory chain complexes required for oxidative phosphorylation (OXPHOS) activity are encoded by both mitochondrial and nuclear genomes, and coordinated expression from both genomes is crucial to allow the stoichiometric assembly and function of these complexes ^37^. Defective import, folding or assembly of these complexes is sensed by mitochondrial protein quality control systems that activate a feedback signaling pathway, dubbed the mitochondrial unfolded protein response (UPR), in order to recover mitochondrial homeostasis ^38^. Similarly, the cytosol and endoplasmic reticulum (ER) are exposed to nascent polypeptides and require dedicated protein-folding machinery. To adjust folding capacity and proteostasis, eukaryotic cells have evolved organelle-specific UPRs ^39^. The common principles of the UPR are the dynamic activation of signal transduction pathways involving transient attenuation of protein synthesis and load and a transcriptional response aimed at increasing organelle capacity for handling unfolded proteins, allowing metabolic adaptation. In cases of prolonged or excessive UPR activation in which homeostasis cannot be re-established, cells are committed to death.

In the present study, for the first time, we demonstrated a role for FANCD2 in the UPR that links mitochondrial dysfunction with genome instability. We show that CFS gene transcription is dependent on mitochondrial activity and is induced by mitochondrial stress and UPR activation. FANCD2 depletion induces mitochondrial dysfunction and CFS gene expression, leading to CFS instability, while attenuation of OXPHOS metabolism decreases CFS gene transcription and rescues chromosome fragility. Conversely, genetic or pharmacological activation of the UPR induces CFS gene transcription and FANCD2 relocalization to CFS genes, dampening the UPR and preventing CFS instability. FANCD2 binding to CFS is dependent on CFS gene transcription and increases in a dose-dependent manner. In addition, we show that FANCD2 is dispensable for maintaining CFS stability in the absence of transcription. Mechanistically, we demonstrate that the UPR-dependent induction of CFS genes is mediated by Ubiquitin-Like Protein 5 (UBL5), which was previously shown to be involved in mitochondrial UPR signaling in *C. elegans*, and that breaking up this signaling partially restores chromosome stability. We propose that CFSs are part of a metabolic checkpoint, and by tuning the UPR with CFS replication, FANCD2 promotes metabolic homeostasis and genome integrity.

## Results

### FANCD2 depletion induces CFS gene expression

Previous studies have shown that transcription of large CFS genes is involved in CFS instability, by either inducing transcription-replication conflicts or modifying their replication dynamics^2, 5^. We thus analyze the impact of FANCD2 on CFS gene expression and stability in a model cell line – HCT116 - in which CFSs have been previously characterized ^40^. Knockdown of FANCD2 increased the transcription of all mapped large CFS genes, whereas the expression of *PTPRG*, a large gene close to *FHIT* in the FRA3B region, was unchanged (Fig. 1a). The increased CFS gene transcription in FANCD2-depleted cells was associated with increased CFS instability, measured by fluorescence in situ hybridization (FISH) (Fig. 1b, c). We verified the upregulation of FHIT expression at the protein level and the increased transcription by measuring nascent *FHIT* RNA transcripts using 5-ethynyl uridine (EU) to label newly synthesized RNA (Fig. 1d). The increased FHIT expression was confirmed using two independent siRNAs targeting FANCD2 (Fig. 1e); the upregulation was also observed after FANCD2 depletion in HeLa and RKO cells and, to a lesser extent, after downregulation of the FANC core protein FANCA (Fig. 1f to h).

**Figure 1:**
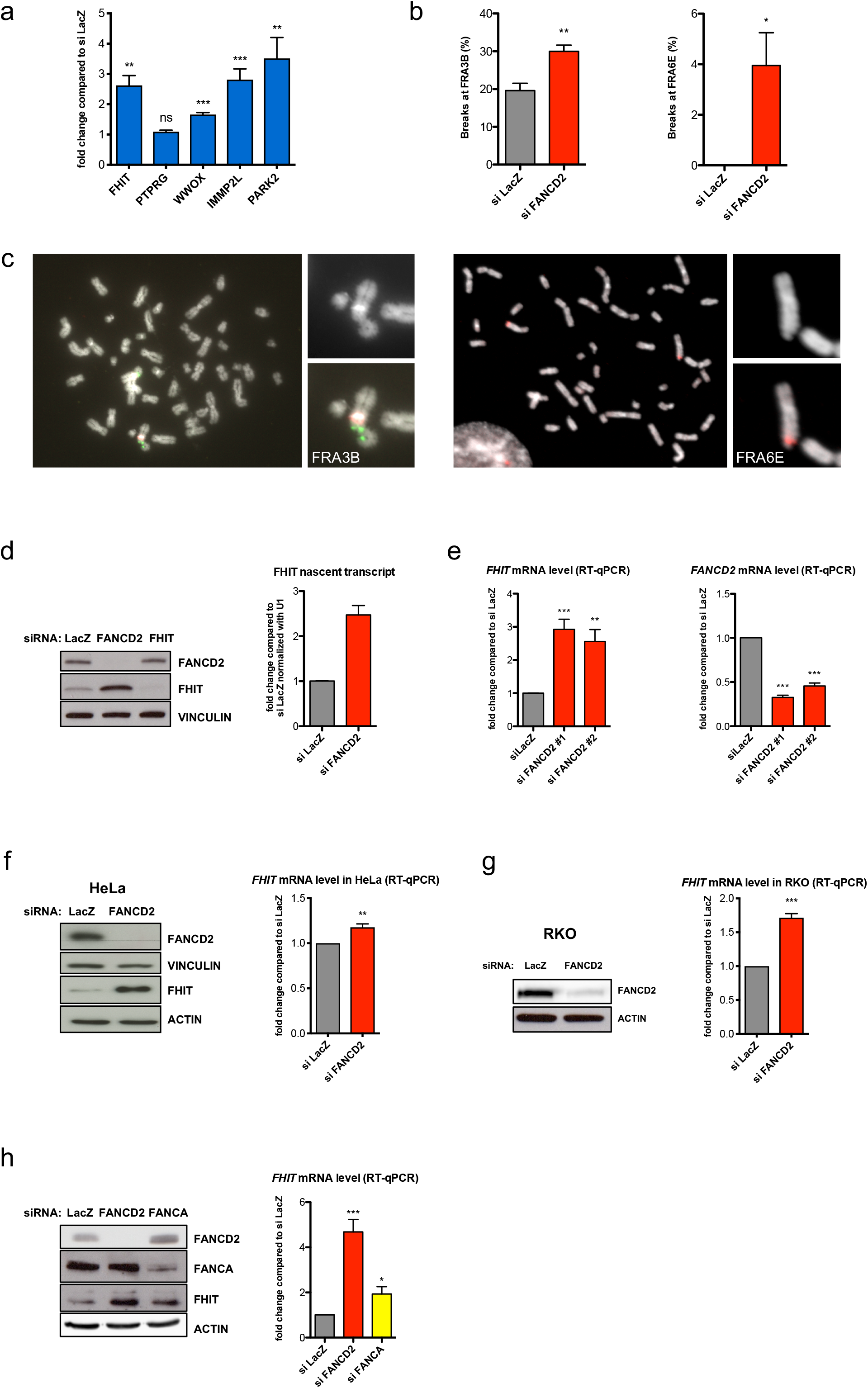
FANCD2 induces CFS gene expression. a) mRNA levels of large CFS genes measured by RT-qPCR after siFANCD2 treatment compared to levels after siLacZ treatment. b) Frequency of FRA3B and FRA6E breakage presented as the percentage of chromosome 3 and chromosome 6 homologs, with breaks at FRA3B and FRA6E, respectively. c) Left, example of FISH analysis of metaphase spread from siFANCD2-transfected cells after treatment with 0.3 µM APH stained for DNA (DAPI, grayscale), a centromeric probe for chromosome 3 (red) and a *FHIT*/FRA3B FISH probe (green). Right, example of FISH analysis of metaphase spread from siFANCD2-transfected cells after treatment with 0.3 µM APH stained for DNA (DAPI, grayscale), and a *PARK2*/FRA6E FISH probe (red). d) Western blot of whole-cell lysate of control, FANCD2, and FHIT siRNA-transfected HCT116 cells showing the increased FHIT protein level in FANCD2-depleted cells (left). Cells were transfected with siRNA against *FHIT* as a specificity control for the FHIT antibody. Quantification of nascent EU-labeled *FHIT* transcripts after control or FANCD2 siRNA transfection by RT-qPCR normalized to *U1* (*RNU1-1*) RNA gene expression (right). e) RT-qPCR analysis of FHIT expression using two independent FANCD2 siRNAs (left). FANCD2 downregulation was estimated by RT-qPCR (right). f) Western blot of whole-cell lysate of control and FANCD2 siRNA-transfected HeLa cells (left). mRNA levels of *FHIT* measured by RT-qPCR after treatment with control or FANCD2 siRNA (right). g) Western blot of whole-cell lysate of control and FANCD2 siRNA-transfected RKO cells (left). Note that RKO cells expressed very low levels of FHIT and that FHIT protein was not detected by Western blot in this cell line. Quantification of *FHIT* mRNA levels by RT-qPCR after treatment of cells with control or FANCD2 siRNA (right). h) FHIT expression increases after FANCD2 and FANCA depletion in HCT116 cells relative to the control, observed at the protein level by Western blot (left) and the mRNA level by RT-qPCR (right).

### FANCD2 binds to CFS genes and prevents their instability in a transcription dependent manner

To decipher the role of transcription in CFS instability and in FANCD2 function, we deleted the *FHIT* promoter in HCT116 cells using the CRISPR/Cas9 system and verified that *FHIT* transcription was suppressed (*FHIT*-KO, Fig. 2a). We then investigated FRA3B instability by analyzing metaphase spreads for the frequency of FRA3B breakage by FISH after treatment with low doses of aphidicolin (APH), which specifically induces CFS instability ^41^. As shown in Fig. 2b, FRA3B breaks were strongly reduced in *FHIT*-KO cells compared to the parental (*FHIT* wt) counterparts, consistent with the role of transcription in inducing CFS instability. Most importantly, residual breaks at FRA3B that formed in control (siLacZ) cells in the absence of *FHIT* transcription (7.82%) were not increased by FANCD2 depletion (4.06%), demonstrating that FANCD2 role at CFSs is linked to the transcription of the corresponding gene. Consequently, we examined whether FANCD2 binding to CFSs was dependent on CFS gene transcription. We performed FANCD2 ChIP followed by qPCR to analyze FANCD2 binding to the *FHIT* gene in wt and *FHIT*-KO cells. Interestingly, FANCD2 binding to *FHIT* was almost abolished in the absence of transcription, whereas binding to other CFS genes was not affected (Fig. 2c). Therefore, FANCD2 is targeted to CFS genes and prevents their fragility in part in a transcription-dependent manner.

**Figure 2:**
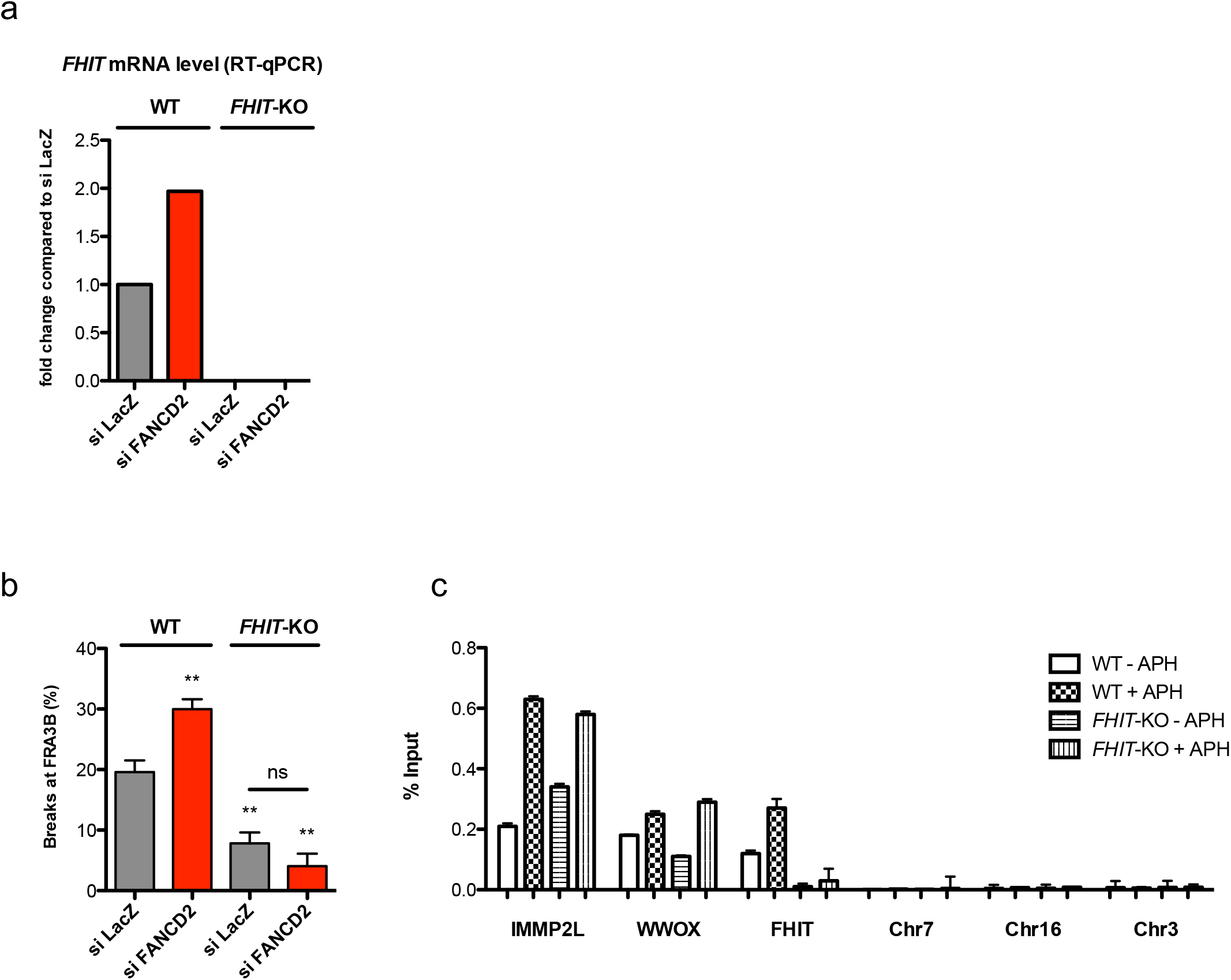
FANCD2 binds to CFS genes and prevents their instability in a transcription dependent manner. a) FHIT mRNA level detected by RT-qPCR in WT and *FHIT*-KO cells after control or FANCD2 siRNA transfection. b) FRA3B instability in WT and *FHIT*-KO cells transfected with control or FANCD2 siRNA and treated with 0.3 µM APH. c) FANCD2 ChIP followed by qPCR in wild-type (WT) and *FHIT*-KO cells treated or not with 0.3 µM APH. The results are expressed as the percentage of input.

### Genome-wide analysis of FANCD2 targets identifies an enrichment for mitochondrial UPR-response elements at FANCD2 binding sites

To identify regulatory sequences modulating CFS gene transcription and FANCD2 function, we analyzed FANCD2 genomic binding sites by chromatin immunoprecipitation sequencing (ChIP-seq) of the endogenous protein in samples that were untreated or had undergone replicative stress induced by low doses of APH, which induces FANCD2 accumulation and persistence to CFS ^18, 20^. In both conditions, FANCD2 peaks were remarkably enriched in genic regions compared with randomly located peaks (Supplementary Fig. 1a). We then analyzed the regions where FANCD2 was most bound after APH treatment (the top induced genes) and observed that they were enriched in large loci (Supplementary Fig. 1b), the vast majority of which corresponded to previously characterized CFSs (Table 1), in agreement with the results of recent studies in human U2OS and chicken DT40 cells ^18, 19^. FANCD2 was also recruited to some CFS genes, such as *FHIT* and *WWOX*, under the untreated condition (Fig. 3a). Moreover, a closer inspection revealed that most of these genes have reported functions in mitochondrial activity, ER dynamics and secretory pathways (Table 1). We then performed a bioinformatic analysis to identify DNA motifs enriched at FANCD2 genomic binding sites. Remarkably, this analysis revealed a significant enrichment of MURE1 and MURE2 elements at FANCD2 binding sites both in the untreated condition and following APH treatment (Fig. 3b). Strikingly, these two elements have been discovered as mitochondrial UPR response elements (MURE), even if the factors binding to the latter sites were not identified ^42, 43^. In addition, we identified recurrent sequences of 54 bp or 63 bp containing combined MURE1, CHOP and MURE2 elements at the promoter and/or in the body of some CFS genes (Fig. 3a and Supplementary Fig. 2). This triplet of elements is a reportedly highly specific functional module required for mitochondrial UPR regulation^43^. Interestingly, we found that FANCD2 enrichment at CFSs increased with gene expression level (Supplementary Fig. 1c). Together, these data suggest an interplay between UPR signaling and FANCD2 in the transcription and stability of CFS genes.

**Table 1:**
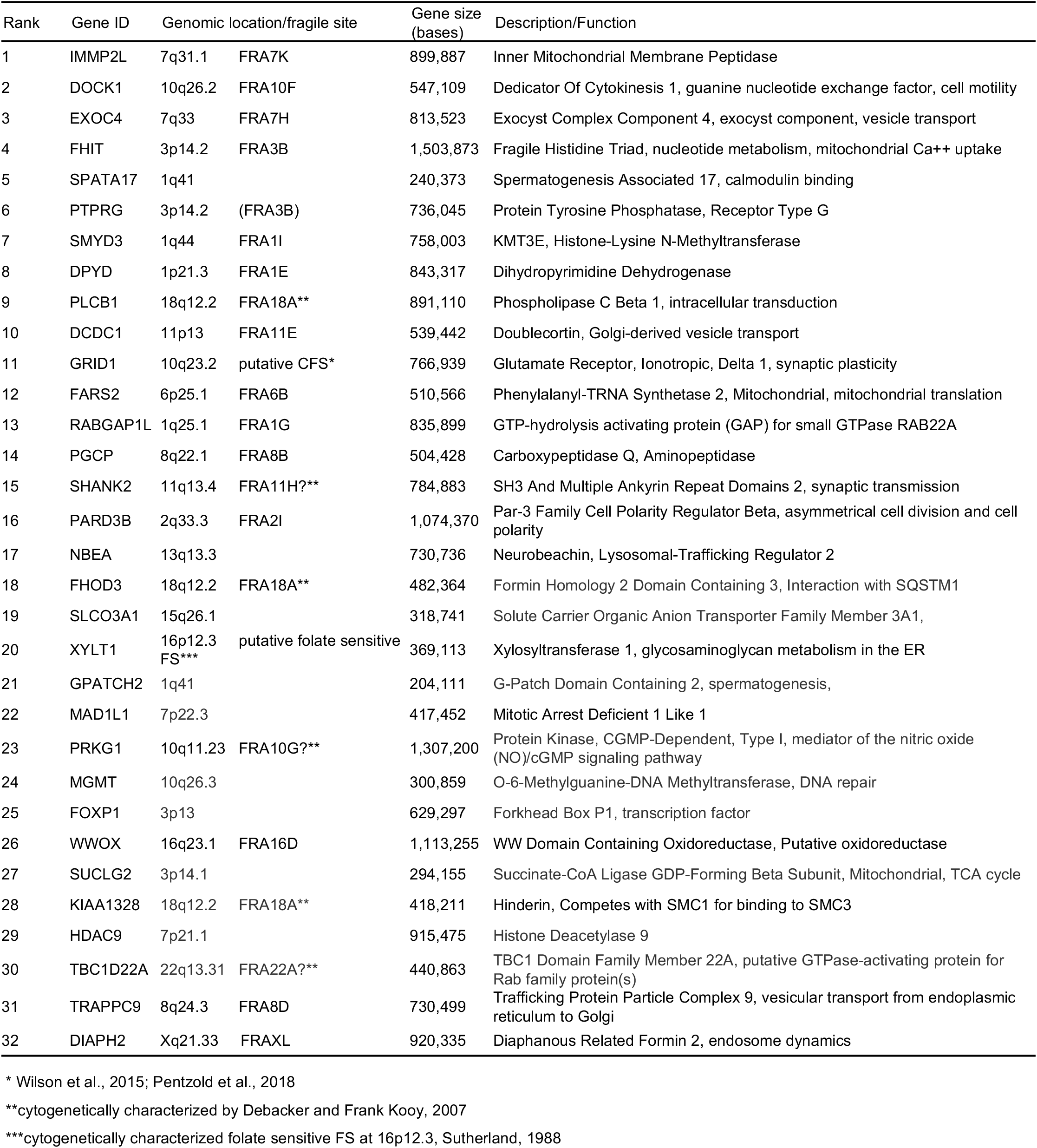
FANCD2 top induced genes

**Figure 3:**
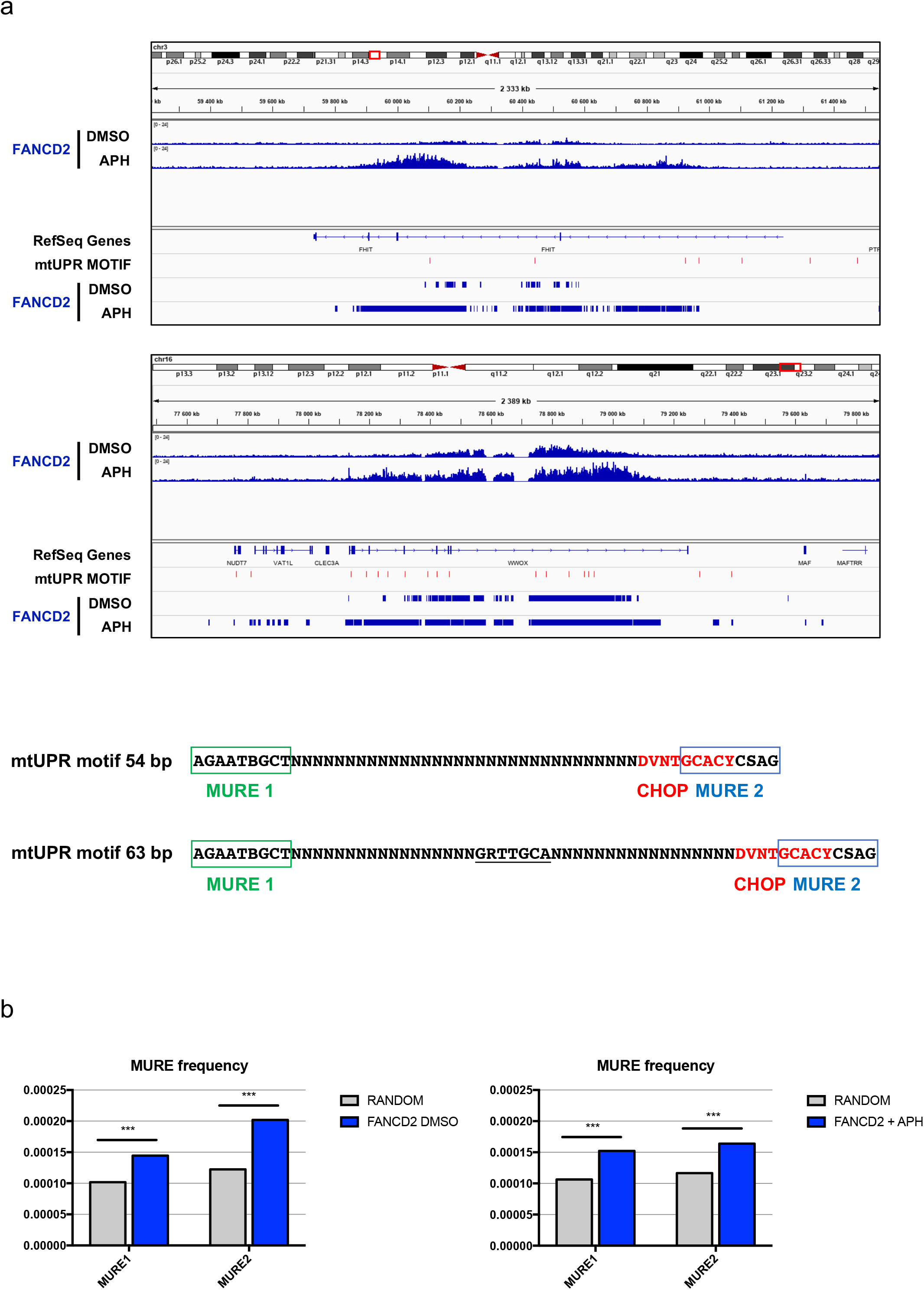
Genome-wide analysis of FANCD2 targets identifies an enrichment for UPR-response elements at FANCD2 binding sites. a) IGV visualization of FANCD2 enrichment along the CFS genes *FHIT* and *WWOX* in the presence or absence of APH and the relative position of mitochondrial UPR (mtUPR) motifs (red tags) and FANCD2 peaks. FANCD2 ChIP seq data were scanned with RSAT (Regulatory Sequence Analysis Tools)-matrix-scan ^66, 67^ to identify instances of MURE1, MURE2 (mitochondrial UPR response elements) and CHOP motifs as they were defined by Aldridge et al (2007) and Munch and Harper (2016) ^42, 43^. mtUPR motifs represent sequences of 54 bp or 63 bp with MURE1-CHOP-MURE2 consensus elements as indicated. The conserved CHOP consensus between MURE1 and MURE2 elements as reported by Aldridge et al. is underlined. Notice that a 10 bp consensus sequence for CHOP described in Munch and Harper is partially overlapping the MURE2 element. b) Frequency of MURE1 and MURE2 elements at FANCD2 binding sites relative to a random control. The matches scored for each motif were compared to those detected in sets of random sequences of identical length from human genome GRCh37-hg19. P values of the probability to obtain a similar score were ***p<1e-16 for each motif.

### UPR triggers CFS gene expression and FANCD2 binding at CFSs

Activation of the mammalian mitochondrial UPR can be triggered by stress in the mitochondria or in the ER ^38^ that communicate through mitochondria-associated ER membrane (MAM) contacts ^44, 45^. To analyze if the UPR is involved in CFS gene transcription, we pharmacologically induced the UPR using either carbonyl cyanide m-chlorophenyl hydrazone (CCCP), a mitochondrial uncoupler, or thapsigargin (TG), a sarco/ER Ca^2+^-ATPase (SERCA) inhibitor and a known inducer of ER stress. Interestingly, both treatments induced the transcription of CFS genes, which was further increased after FANCD2 depletion, indicating that CFS genes respond to UPR activation and that FANCD2 dampens this response (Fig. 4a).

**Figure 4:**
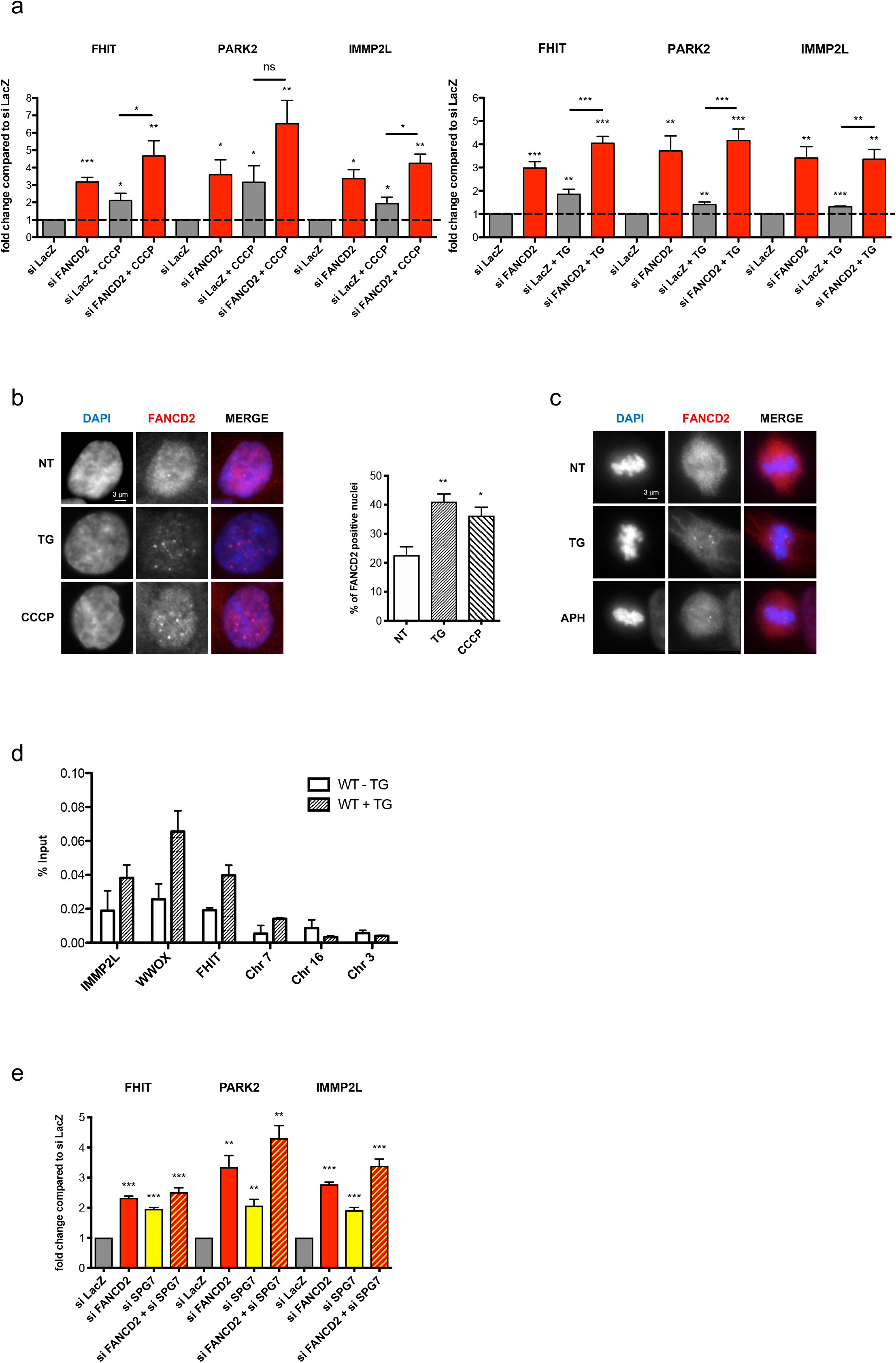
UPR triggers CFS gene expression and FANCD2 binding at CFSs. a) Left, RT-qPCR-based analysis of CFS gene expression in siLacZ- and siFANCD2-transfected cells treated or not with 10 µM CCCP for 8 hr. Right, RT-qPCR-based analysis of CFS gene expression in siLacZ- and siFANCD2-transfected cells treated or not with 1 mM TG for 8 hr. b) Left, examples of immunofluorescence staining of FANCD2 in cells treated with TG or CCCP or in cells that were not treated (NT); FANCD2 is shown in red, and DNA (DAPI) is in blue in the merged image. Right, percentage of FANCD2-positive nuclei; FANCD2 foci were counted in at least 50 nuclei in each of four different experiments, and nuclei with more than 5 spots were quantified. c) Immunofluorescence staining of FANCD2 in metaphase cells treated or NT with TG or APH. d) FANCD2 ChIP followed by qPCR in cells treated or not with 1 mM TG. The results are expressed as the percentage of the input. Chromosome 7 (Chr 7), Chromosome 16 (Chr 16), and Chromosome 3 (Chr 3) were used as control regions close to the *IMMP2L*, *WWOX*, and *FHIT* genes, respectively. e) Expression of large CFS genes measured by RT-qPCR after control, FANCD2, SPG7, or FANCD2 and SPG7 siRNA transfection.

In parallel, we looked at FANCD2 behavior after UPR induction by TG or CCCP treatment. UPR activation induced FANCD2 relocalization in nuclear foci, some of which persisted in mitosis, similar to what was observed after APH treatment and suggesting that they correspond to CFSs (Fig. 4b, c). To support this finding, we performed FANCD2 ChIP followed by qPCR after UPR induction. We detected a specific enrichment of FANCD2 at CFS genes (Fig. 4d), showing that UPR signaling promotes the recruitment of FANCD2 to CFSs.

We then used a genetic strategy to specifically perturb mitochondrial proteostasis. We downregulated spastic paraplegia 7 (*SPG7*), the gene encoding the paraplegin matrix AAA peptidase subunit, a mitochondrially localized, membrane associated protease. SPG7 downregulation increases the load of unfolded proteins in the mitochondria, activating the mitochondrial UPR uncoupled from accumulation of reactive oxygen species ^46^. As shown in Fig. 4e, SPG7 depletion by RNAi increased CFS gene transcription, similar to what is observed after FANCD2 downregulation, demonstrating that CFS genes expression is triggered by a mitochondrial-dependent stress signaling and that FANCD2 counteracts it.

### Attenuation of mitochondrial respiration or UBL5 depletion mitigates CFS instability in FANCD2-depleted cells

To unravel the role of mitochondrial stress and UPR signaling in CFS instability, we examined the consequences of FANCD2 deficiency on mitochondrial function. Downregulation of FANCD2 elicited a specific defect in electron transport between complexes I and III of the respiratory chain, leading to increased oxygen consumption and decreases in ATP synthesis and the ATP/AMP ratio (Fig. 5a). The impaired mitochondrial energy production and the decrease in the ATP/AMP ratio was accompanied by an increased lactate dehydrogenase (LDH) activity at time points after FANCD2 depletion, suggesting a shift to glycolytic metabolism to compensate for the OXPHOS defect (Supplementary Fig. 3). We then tested whether the expression of CFS genes was dependent on mitochondrial OXPHOS activity using sodium azide (NaN_3_) to inhibit mitochondrial respiration. Treatment with NaN_3_ decreased the expression of all tested CFS genes in both control and FANCD2-depleted cells (Fig. 5b). Finally, we ascertained whether physiological attenuation of OXPHOS metabolism attenuates CFS gene expression and instability by culturing cells at low oxygen tension (3%) and measuring CFS transcription. Compared to cells cultured in 20% O_2_, FANCD2-depleted cells cultured in 3% O_2_ showed a significant reduction in the transcription of CFS genes, except for *PARK2* that was upregulated under low oxygen concentration (Fig. 5c); *PARK2* expression may be induced to promote the shift to glycolytic or fatty acid metabolism ^47–49^. We then assessed whether chromosome breakage is decreased at 3% O_2_ after depletion of FANCD2. Remarkably, the global frequency of breaks in metaphase chromosomes was significantly rescued in FANCD2-depleted cells cultured in 3% O_2_ compared with cells cultured in 20% O_2_ (Fig. 5d), even if a low frequency of breaks specifically occurred at *PARK2*/FRA6E (Fig. 5e), showing a striking correlation between increased CFS gene transcription and instability. Taken together, these data indicate that CFS gene expression is linked to mitochondrial activity and is associated with mitochondrial dysfunction induced after FANCD2 depletion.

**Figure 5:**
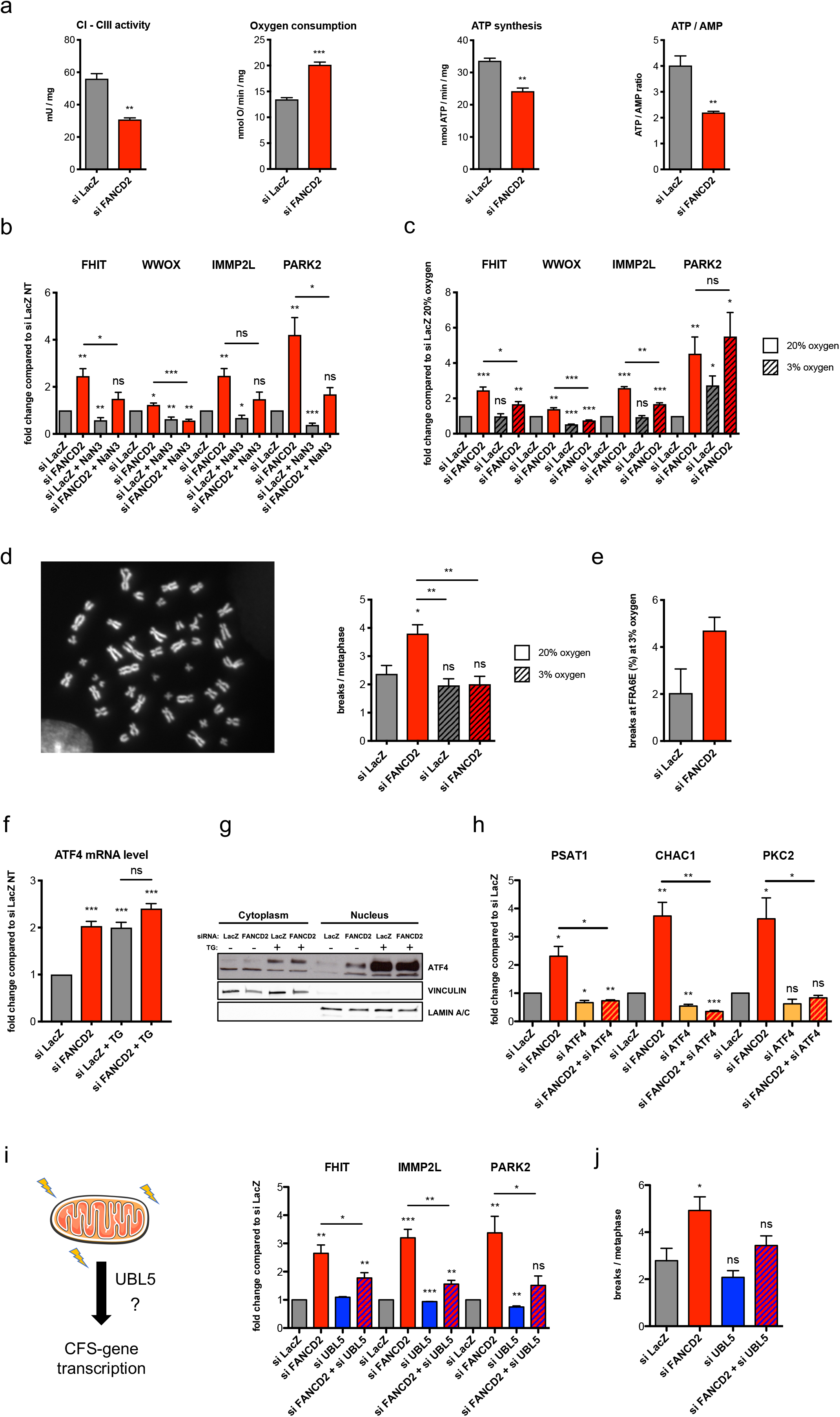
UBL-5 depletion and attenuation of mitochondrial respiration mitigates CFS instability in FANCD2-depleted cells. a) Parameters for mitochondrial activity and cellular energy status analyzed after control or FANCD2 siRNA transfection. b) RT-qPCR-based analysis of CFS gene expression in cells after control or FANCD2 siRNA transfection and treated or not with 20 mM NaN3 for 10 hr. c) RT-qPCR analysis of CFS gene expression after control or FANCD2 siRNA transfection of cells maintained at 3 or 20% oxygen. d) Chromosome fragility in cells transfected with control or FANCD2 siRNA, treated with 0.3 µM APH and maintained at 3 or 20% oxygen. Left, example of a DAPI-stained metaphase spread; the arrows indicate a break. Right, total breaks are scored as the mean number of breaks per metaphase. At least 50 metaphases were analyzed per condition and experiment (n=3). e) Frequency of FRA6E breaks in cells maintained at 3% oxygen, transfected with control or FANCD2 siRNA and treated with 0.3 µM APH. f) ATF4 expression measured by RT-qPCR in siLacZ- and siFANCD2-transfected cells, treated or not with 1 mM TG. g) Western Blot detection of ATF4 in cytoplasmic and nuclear fractions of siLacZ- or siFANCD2-transfected cells treated (+) or not (-) with 1 mM TG. Vinculin and lamin A/C were used as loading controls for cytoplasmic and nuclear fractions, respectively. h) Expression of ATF4 target genes measured by RT-qPCR after control, FANCD2, ATF4, or FANCD2 and ATF4 siRNA transfection. i) Left, graphical representation of mitochondrial UPR activation of CFS genes. Right, expression of large CFS genes measured by RT-qPCR after control, FANCD2, UBL5, or FANCD2 and UBL5 siRNA transfection. j) Chromosome fragility in cells transfected with control, FANCD2, UBL5, or FANCD2 and UBL5 siRNA and treated with 0.3 µM APH, scored as the mean number of breaks per metaphase. At least 50 metaphases were analyzed per condition and experiment (n=4).

We then investigated the mechanism underlying UPR signaling and its role in FANCD2 function and CFS stability. ATF4 is a transcription factor activated upon UPR induction and a key effector of mitochondrial stress response in mammalian cells^50^. Remarkably, FANCD2 downregulation induced the transcription of ATF4, as well as its nuclear relocalization and the activation of its targets *CHAC1*, *PCK2*, and *PSAT1* (Fig. 5f to h), showing the activation of mitochondrial stress signaling. Consistently, ATF4 knockdown decreased the expression of these genes and abrogated their induction after FANCD2 depletion (Figure 5d). However, ATF4 depletion did not restore the increased CFS gene transcription observed after FANCD2 depletion (Supplementary Fig. 4), suggesting that CFS genes may be regulated by FANCD2 in an ATF4-independent manner. UBL5 is a ubiquitin-like protein involved in mitochondrial UPR in *C. elegans* ^51^, that has been shown to promote the FANC pathway functionality. Strikingly, UBL5 knockdown significantly reduced the upregulation of CFS genes in FANCD2-depleted cells (Fig. 2i), revealing the functional interplay between FANCD2 and UBL5 in regulating the mitochondrial UPR in human cells. Remarkably, reducing UPR signaling by downregulating UBL5 in FANCD2-depleted cells decreased the chromosome instability after APH treatment (Fig. 2j) at levels similar to control values, emphasizing that suppression of mitochondrial stress signaling mitigates chromosome fragility in the absence of a functional FANC pathway.

Collectively, these data highlight a new role of FANCD2 as a regulatory component of the UPR involved in mito-nuclear communication, counteracting mitochondrial stress and attuning UPR-mediated CFS gene transcription to prevent replication stress and CFS instability (Fig. 6).

**Figure 6:**
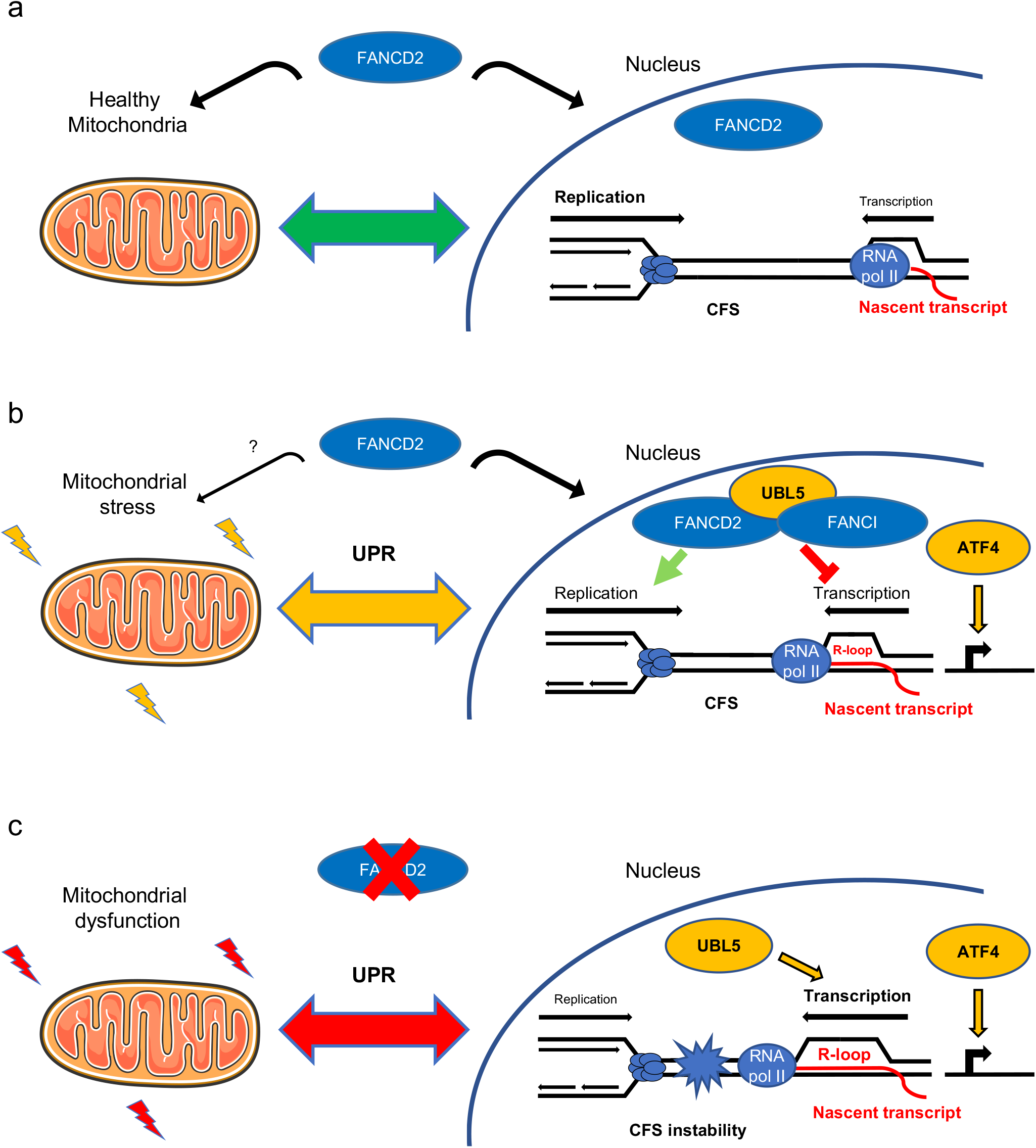
FANCD2 maintains mitochondrial homeostasis and genome stability by tuning the UPR. a) FANCD2 coordinates nuclear and mitochondrial activities to prevent mitochondrial dysfunction and maintain the UPR in check. b) Upon mitochondrial stress, FANCD2 relocalizes to CFS and dampens the UPR, limiting transcription-replication conflicts. c) In the absence of FANCD2, mitochondrial dysfunction and UPR-induced, UBL5-dependent CFS gene transcription leads to transcription-replication collisions and CFS instability.

## Discussion

The coordinated regulation of mitochondrial and nuclear activities is essential for cellular function and metabolic homeostasis. Mitochondria are signaling hubs whose activity is monitored through metabolic readouts produced during oxidative metabolism, such as the levels of metabolites, nucleotides, reactive oxygen species, the rate of ATP production or the level of misfolded proteins ^52^. In recent years, several studies have highlighted the existence of retrograde pathways, called UPRs, that signal organelle- and compartment-specific stress and activate nuclear programs to adjust cellular metabolic activities and recover homeostasis. The mitochondrial UPR has been primarily characterized in *C. elegans*, but some of the components linking mitochondrial protein misfolding to the nucleus have been recently identified in mammalian cells ^53^. In this study, we show for the first time that FANCD2 participates in this pathway in human cells, revealing a new player in this mitochondrial nuclear crosstalk. FANC proteins are best known for their role in the maintenance of genome stability. However, several studies have shown that they also perform non-canonical functions in mitochondria ^28, 54^. In this study, we identified a dual function of FANCD2 in counteracting mitochondrial stress and UPR activation and in dampening the UPR-induced transcription of CFS genes. Large CFS genes may be exquisitely sensitive rheostats of cellular metabolic activity. For example, variations in dNTP biosynthesis and ROS have been shown to directly modulate replisome architecture and replication fork velocity ^55^. CFSs are late replicating, and slowing their replication further increases the risk of incomplete replication and breakage at the time of mitosis ^6^. Furthermore, the timing of CFS replication is modulated by transcription ^5^, and the failure to coordinate these two processes leads to CFS breakage ^2^. Recent work has highlighted the role of the FANC pathway in coordinating replication and transcription, and FANCD2 has been shown to enable efficient replication of CFSs by preventing or resolving R-loop formation ^23–25^. In the present study, we revealed a new layer of CFS regulation that links their transcription to mitochondrial activity through the mitochondrial UPR. We show that CFSs are regulated by an UPR arm that is separate from ATF4, a key regulator of the mitochondrial retrograde response^50^, and is modulated by a UBL5-FANCD2 axis. In this branch, UBL5 acts as an activator and FANCD2 as a suppressor, which may constitute a feedback loop for fine tuning the UPR. Indeed, it has been reported that UBL5 stabilizes FANCI and FANCD2 and promotes their interaction ^56^, which may titrate UBL5 leading to UPR attenuation. Further work will be required to elucidate the dynamics of this novel important pathway that mediates UPR regulation. Depletion of FANCD2 induces mitochondrial stress and activates the ATF4 pathway, which rewires mitochondrial metabolism ^50, 57, 58^ and may be beneficial, at least in part, during the recovery from stress. It is likely that transient activation of CFS genes is also required to recover mitochondrial or ER homeostasis ^44^. In this scenario, mild mitochondrial dysfunction and UPR activation would be beneficial and constitute a feedback mechanism to ensure cellular homeostasis. However, prolonged or excessive mitochondrial dysfunction and UPR activation would lead to replication stress and conflicts between replication and transcription (Fig. 6). Noteworthy, it has been reported that TG-mediated UPR induction reduces replication fork progression and origin firing^59^.

We demonstrated that transcription is required for FANCD2 recruitment to CFS genes and for its function in CFS maintenance. In addition, we found that FANCD2 enrichment at CFSs is proportional to the level of CFS gene transcription (Supplementary Fig. 1b) and is promoted both by UPR induction (Fig. 4e, f) and by replication stress (Fig. 3b and Supplementary Fig. 2). Therefore, we propose that CFS loci behave as both cis- and trans-acting components of the UPR that become unstable above a threshold of UPR activation and replication stress. The encounters between replication and transcription and R-loop formation would generate the substrate for FANCD2 binding and/or retention at CFSs and trigger the activation of the FANC pathway ^18, 23, 24, 60^, constituting a metabolic and genome surveillance checkpoint.

Interestingly, we have found that FANCD2 binding sites are enriched in MURE elements, and that CFS genes contain a variable number of mitochondrial UPR modules of 54 bp or 63 bp (Fig. 3c and Supplementary Fig. 2). Since both MURE1 and MURE2 elements and the combination of MURE1-CHOP-MURE2 motifs have been shown to be involved in mitochondrial UPR regulation ^42, 43^, it is tempting to speculate that CNVs and rearrangements targeting CFSs may affect the magnitude of the CFS gene response to UPR and, as a consequence, cellular stress resistance and metabolic adaptation ^61^ during normal development and in cancer ^62^. Intriguingly, the UPR has also been shown to be a major metabolic checkpoint that regulates hematopoietic stem cell function and integrity ^63–65^. It will be crucial in the future to better characterize how the FA pathway regulates the UPR during hematopoiesis and whether it modulates the expression and stability of specific CFSs in hematopoietic cells.

## Materials and Methods

### Cell culture

The HCT116 cell line was maintained in McCoy’s 5A medium (ATCC), and RKO and HeLa cells were maintained in Dulbecco’s modified Eagle’s medium (DMEM) (Gibco) at 37°C in a humidified atmosphere under 5% CO_2_. Media were supplemented with 10% fetal bovine serum (FBS), 1 mM sodium pyruvate, 100 U/mL penicillin and 100 µg/mL streptomycin. All cell lines were purchased from ATCC. HCT116 FHIT-KO cells were generated by CRISPR/Cas9 genome editing of parental HCT116 cells. For experiments requiring low oxygen conditions, cells were maintained in an incubator with an O_2_ control system (HERAcell 150i, Thermo Scientific).

### siRNA transfection

siRNA duplex oligonucleotides were purchased from Ambion to target FHIT (#AM16708) and from Eurogentec to target the other assayed genes. The siRNA sequences are provided in Supplementary Table S1. For all siRNA experiments, cells were transfected with siRNAs at a final concentration of 20 nM using INTERFERin (Polyplus) according to the manufacturer’s instructions. Following siRNA transfection, knockdown of gene expression was assessed by Western blot or qRT-PCR analysis. Unless otherwise indicated, cells were collected for total cell lysate preparation, subcellular fractionation, biochemical assays, and qRT-PCR analysis 48 hr after transfection.

### Western blotting and subcellular fractionation

For total lysates, cells were disrupted in lysis buffer (50 mM Tris-HCl, 20 mM NaCl, 1 mM MgCl_2_, and 0.1% SDS) containing a protease and phosphatase inhibitor cocktail (Roche) supplemented with 0.1% endonuclease (benzonase, Millipore) for 10 min at room temperature with rotation. For fractionation analysis, cells were lysed using an NE-PER kit (ThermoFisher) following the manufacturer’s instructions. Laemmli buffer containing beta-mercaptoethanol was added to the samples, which were subsequently boiled for 5 min at 95°C. The proteins were separated on SDS-PAGE denaturing gels (Bio-Rad) and transferred to nitrocellulose membranes (Bio-Rad). Next, the membranes were blocked with phosphate-buffered saline (PBS)-milk (5%) or PBS-bovine serum albumin (BSA) (3%) for 1 hr, and signals were visualized using WesternBright ECL (Advansta) on a digital imaging system (GeneGnome, Syngene) or using Amersham Hyperfilm ECL film (GE) on a table-top processor (Curix 60, AGFA). The antibodies used in this study are listed in Supplementary Table S1.

### Quantitative RT-PCR

Total cellular RNA was extracted with the ReliaPrep RNA Cell Miniprep System (Promega), and 1 µg of RNA was used to synthesize cDNA with a High-Capacity RNA-to-cDNA kit (ThermoFisher). PCR primers were purchased from Eurogentec and used in PCRs with SYBR Green Master Mix (ThermoFisher) on a QuantStudio 7 Flex instrument (Applied Biosystems). Relative gene expression was calculated using the ΔΔCq method and normalized to *GAPDH* expression. Values are represented as the fold change compared to the control transfection values (siLacZ). Primer sequences are available in Supplementary Table S1.

### Cell treatments and chemicals

Replicative stress was induced by treating cells with 0.3 µM APH (Sigma A0781) for 20 hr. The UPR was induced by treatment with 1 mM TG (Interchim 42759J) or 10 µM CCCP (Sigma C2759) for 8 hr. For OXPHOS inhibition, sodium azide (NaN_3_, Sigma S8032) was used at 20 mM for 10 hr.

### Immunofluorescence

Cells grown on glass cover slips were fixed in 4% formaldehyde for 15 min before permeabilization with 0.5% Triton for 10 min at room temperature. After blocking with 3% BSA in PBS containing 0.05% Tween 20, cells were stained overnight with the primary antibody against FANCD2 and then with a secondary antibody, anti-rabbit Alexa Fluor 594 (Invitrogen), for 1 hr at room temperature. Slides were mounted in DAKO mounting medium containing 4’,6-diamidino-2-phenylindole (DAPI) (SouthernBiotech) and examined at a 63× magnification using an epifluorescence microscope (Zeiss Axio Observer Z1) equipped with an ORCA-ER camera (Hamamatsu). The microscope and camera parameters were set for each series of experiments to avoid signal saturation. Image processing and analysis were performed using ImageJ.

### Nascent transcript analysis

Nascent transcripts were captured and analyzed using a Click-iT Nascent RNA Capture Kit from ThermoFisher following the manufacturer’s instructions. Briefly, cells were seeded in a 6-well plate and transfected the next day. Next, 48 hr after transfection, cells were incubated with 0.5 mM EU for 1 hr and harvested for RNA extraction. Then, 5 µg of RNA was biotinylated with 0.5 mM biotin azide and precipitated. Finally, 1 µg of biotinylated RNA was bound to 50 µL of streptavidin magnetic beads and used for cDNA synthesis and qPCR.

### Metaphase spread preparation and FISH analysis

Forty-eight hours after transfection, cells were incubated with or without 0.3 µM APH for 16 hr. Subsequently, the cells were exposed to 100 ng/mL colcemid (Roche) for 3 hr, treated with hypotonic solution (0.075 M KCl) for 15 min and fixed with 3:1 ethanol/acetic acid overnight at −20°C. The cells were then transferred onto slides and dried for one day. For FRA3B analysis, two FISH probes were used: the ZytoLight SPEC FHIT/CEN 3 dual color probe (ZytoLight) and labeled bacterial artificial chromosomes (BACs). Briefly, bacterial strains containing the BACs (RP11-170K19 and RP11-495E23) were grown overnight at 37°C with 12.5 µg of chloramphenicol and extracted using a BACMAX DNA purification kit (Epicentre). Then, DNA was sonicated to obtain fragments shorter than 400 bp, which were then labeled green or red using a PlatinumBright labeling kit (Kreatech) following the manufacturer’s instructions. A PARK2 FISH probe (Empire Genomics) was used for FRA6E analysis. Briefly, slides were sequentially incubated in 70, 90 and 100% ethanol for 2 min and then dried. Subsequently, 10 µL of each probe was added to the slides, and a cover slip was added and adhered with rubber cement on its edges to avoid dehydration. The slides were placed on an automatic hybridizer (Hybridizer, Dako) and heated at 72°C for 2 min and then at 37°C for at least 16 hr. Afterwards, the coverslips were removed in wash buffer (0.5× SSC and 0.1% SDS) at 37°C, and the slides were incubated in wash buffer for 5 min at 65°C to remove nonspecific signals. Finally, the slides were washed with PBS, and DAPI was added with mounting medium for microscopic analysis.

### Chromatin immunoprecipitation and next-generation sequencing (ChIP-seq)

ChIP-seq experiments were performed using Active Motif ChIP sequencing services. First, 1 × 10^7^ HCT116 cells that had been treated with or without 0.3 µM APH were fixed in 11% formaldehyde for 15 min. After cell lysis, 30 µg of chromatin was used for immunoprecipitation using a FANCD2 antibody (Novus). Immunoprecipitated and input DNA were sequenced by Illumina sequencing, which generated 75-nt sequence reads. More than 30 × 10^6^ reads per condition were obtained, and a spike-in adjusted normalization method was applied. The peaks were called using the SICER algorithm and aligned to the human genome build hg19. Integrative Genomics Viewer (IGV) was used to visualize peaks from the genome.

### ChIP and quantitative PCR

After preclearing with magnetic beads for 1 hr, the chromatin from an equivalent of 1 × 10^7^ HCT116 cells was used for immunoprecipitation with a FANCD2 antibody (Novus) or immunoglobulin G as a control. After an overnight incubation at 4°C, the beads were washed and eluted in buffer E (25 mM Tris-HCl [pH 7.5], 5 mM EDTA, and 0.5% SDS), and crosslinking was reversed at 65°C with proteinase K for 6 hr. The DNA was then purified using a QIAquick PCR purification kit (QIAGEN) and eluted in 100 μl of distilled water. The PCR primer pairs are listed in Supplementary Table S1.

### Oxygen consumption measurements

Oxygen consumption was measured at 25°C in a closed chamber using an amperometric electrode (Unisense Microrespiration, Unisense A/S, Denmark). Cells were permeabilized with 0.03 mg/ml digitonin for 1 min, centrifuged for 9 min at 1000 × g and resuspended in a buffer containing 137 mM NaCl, 5 mM KCl, 0.7 mM KH_2_PO_4_, 25 mM Tris-HCl (pH 7.4), and 25 mg/ml ampicillin (Ravera et al., 2013). The same solution was used in the oxymetric measurements. For each experiment, 500,000 cells were used. Finally, 10 mM pyruvate and 5 mM malate were added to stimulate the electron transfer pathway by complexes I, III and IV.

### Electron transfer from complex I to complex III

Electron transfer from complex I to complex III was studied spectrophotometrically by following the reduction of cytochrome *c* at 550 nm. The molar extinction coefficient used for reduced cytochrome *c* was 1 mM^−1^ cm^−1^. For each assay, 50 µg of total protein was used. The assay medium contained 100 mM Tris-HCl (pH 7.4) and 0.03% cytochrome *c*. The reaction was initiated with the addition of 0.7 mM NADH. If electron transport between complexes I and III is conserved, the electrons pass from NADH to complex I, then to complex III via coenzyme Q, and finally to cytochrome *c*.

### ATP and AMP quantification

ATP and AMP were measured according to the enzyme coupling method of Bergmeyer et al. (Bergmeyer HU, Grassl M, Walter HE (1983) Methods of Enzymatic Analysis, Verlag-Chemie, Weinheim, p. 249). For ATP assays, the medium contained 20 µg of sample, 50 mM Tris-HCl pH 8.0, 1 mM NADP, 10 mM MgCl_2_, and 5 mM glucose in a final volume of 1 ml. Samples were analyzed spectrophotometrically before and after the addition of 4 µg of purified hexokinase/glucose-6-phosphate dehydrogenase (Boehringer). The decrease in absorbance at 340 nm due to NADPH formation was proportional to the ATP concentration.

For AMP assays, the medium contained 20 µg of sample, 50 mM Tris-HCl (pH 8.0), 1 mM NADH, 10 mM MgCl_2_, 10 mM phosphoenolpyruvate (PEP), and 2 mM ATP in a final volume of 1 ml. Samples were analyzed spectrophotometrically before and after the addition of 4 µg of purified pyruvate kinase/LDH (Boehringer). The decrease in absorbance at 340 nm due to NADH oxidation was proportional to the AMP concentration. For all biochemical experiments, protein concentrations were determined using the Bradford method.

### Lactate dehydrogenase activity assay

Lactate dehydrogenase (LDH; EC 1.1.1.27) activity was measured to quantify the anaerobic metabolism. The reaction mixtures contained 100 mM Tris-HCl (pH 9), 5 mM pyruvate, 40 μM rotenone and 0.2 mM NADH, with LDH activity expressed as IU/mg of total protein (micromoles/min/mg of protein).

### Fo-F1 ATP synthase activity assay

Evaluation of the Fo-F1 ATP synthase activity was performed as previously described. Briefly, 200,000 cells were incubated for 10 min in a medium containing 10 mM Tris-HCl (pH 7.4), 100 mM KCl, 5 mM KH_2_PO_4_, 1 mM EGTA, 2.5 mM EDTA, 5 mM MgCl_2_, 0.6 mM ouabain and 25 mg/ml ampicillin, and 10 mM pyruvate plus 5 mM malate to stimulate the pathway composed by complexes I, III and IV. ATP synthesis was induced by the addition of 0.1 mM ADP. The reaction was monitored every 30 seconds for 2 min using a luminometer (GloMax® 20/20n Luminometer, Promega Italia, Milan, Italy) for the luciferin/luciferase chemiluminescent method, with ATP standard solutions used at concentrations between 10^−8^ and 10^−5^ M (luciferin/luciferase ATP bioluminescence assay kit CLSII, Roche, Basel, Switzerland). Data are expressed as nmol ATP produced/min/10^6^ cells.

### Statistical analysis

All quantitative data are presented as the means ± SD of at least three independent experiments. Significance was tested using a two-tailed Student’s t-test. Statistical tests were performed using Prism (GraphPad software). P values are indicated as *p≤0.05, **p≤0.01, and ***p≤0.001, with ns indicating not significant (p>0.05).

## Author contributions

P. F. performed research, interpreted data, and wrote the manuscript. B.M. and C.S-R. performed the ChIP-seq analysis and ChIP-qPCR experiments. S.R. and E.C. performed biochemical and metabolic analyses. V. Nähse generated the FHIT-KO cell line. V Naim conceived and supervised the project, carried out research, interpreted data, and wrote the manuscript, with input from other authors.

## Acknowledgements

The authors are grateful to the members of UMR8200 CNRS for helpful discussions and advice, to M. Debatisse for the FHIT-KO cell line, and to M. Debatisse and F. Rosselli for critical reading of the manuscript. The V. Naim laboratory was supported by a European Research Council Starting Grant (ERC-2014-StG-638898 “FAtoUnFRAGILITY”) and a subvention from GEFLUC Paris Ile-de-France. The work of B.M. and C.S-R. was supported by the Fondation pour la Recherche Médicale (AJE20151234749), the Institut National du Cancer-Plan Cancer (ASC15018KSA) and Labex “Who am I?” (ANR-11-LABX-0071 and ANR-11-IDEX-005-02). E.C. is indebted to AIRFA for its support in the activity of the Clinical & Experimental Hematology Unit of G. Gaslini Institute. P.F. was the recipient of a PhD fellowship from University Paris-Sud.

---

**Supplementary Fig. 1.**
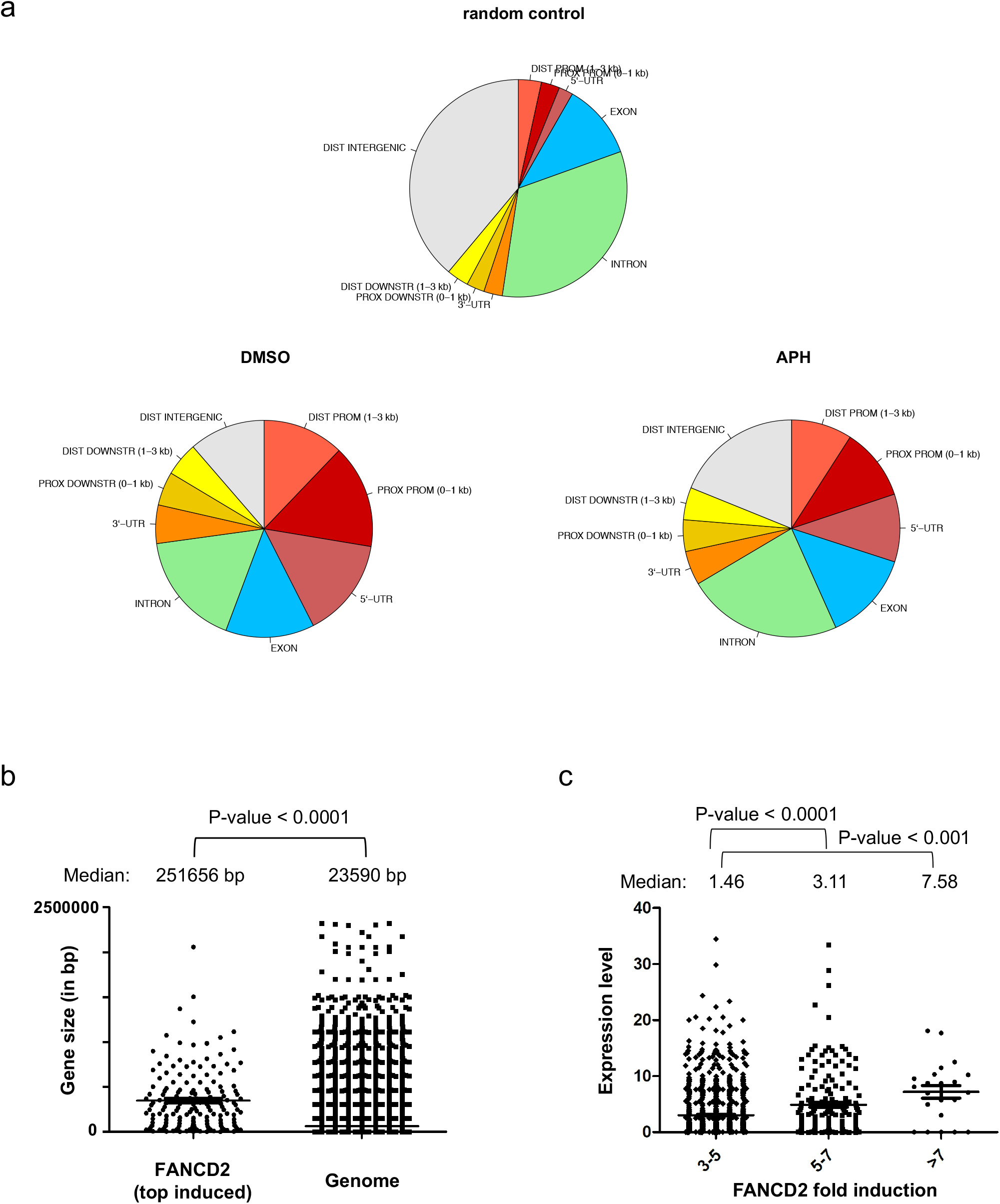
a) FANCD2 location peaks relative to genomic annotations are represented in a pie chart. As a control, randomly located “peaks” were also run against the same genomic features database. b) Comparison of the median gene size between FANCD2 top-induced genes and all genes present in the genome. c) FANCD2 binding fold induction according to the gene expression level. Gene expression data were obtained from ENCODE: Experimental Summary ENCSR000CWM. Gene expression values are in log2 FPKM (<u>F</u>ragments <u>P</u>er <u>K</u>ilobase of exon per <u>M</u>illion reads) units.

**Supplementary Fig. 2.**
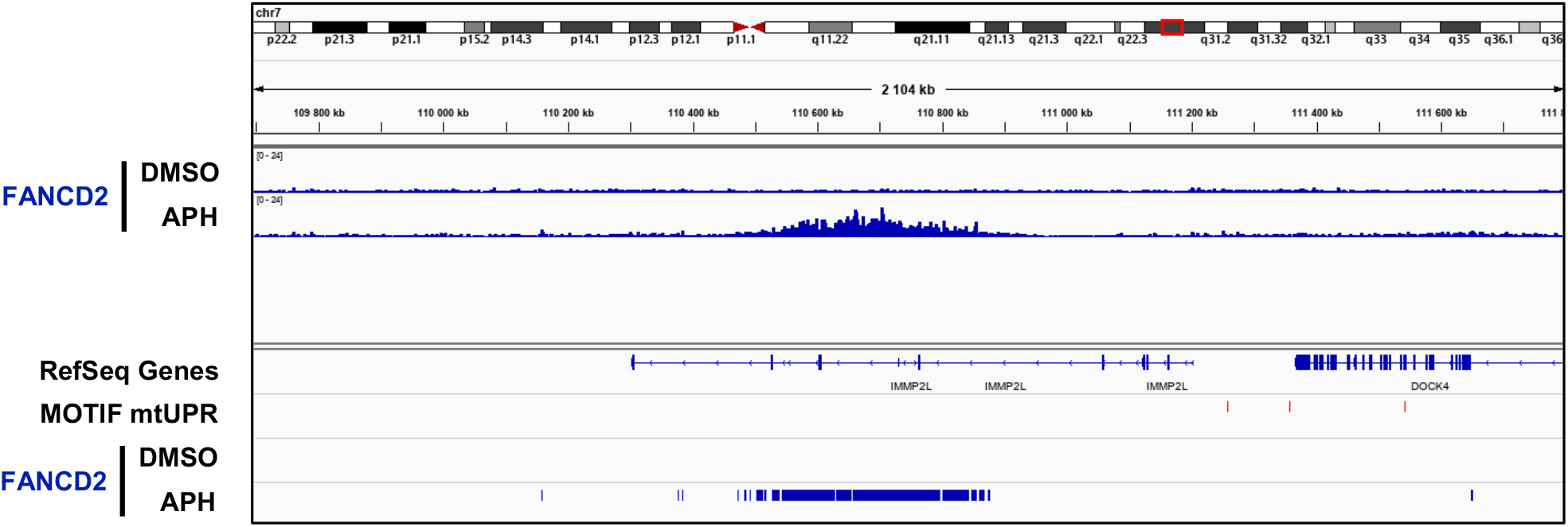
IGV visualization of FANCD2 enrichment along the CFS gene *IMMP2L* in the presence or absence of APH and location of mitochondrial UPR (mtUPR) motifs (red tags) and FANCD2 peaks.

**Supplementary Fig. 3.**
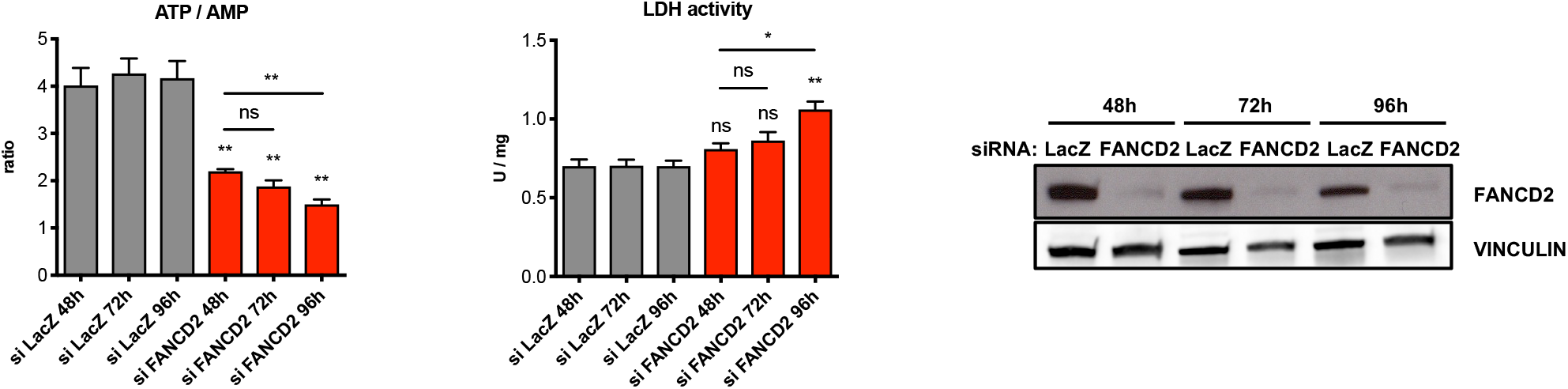
ATP/AMP ratio and LDH activity measured at 48, 72 and 96 hr after transfection with control or FANCD2 siRNA (left). Western blot of whole-cell lysate of control and FANCD2 siRNA-transfected cells at the indicated time points after transfection (right).

**Supplementary Fig. 4.**
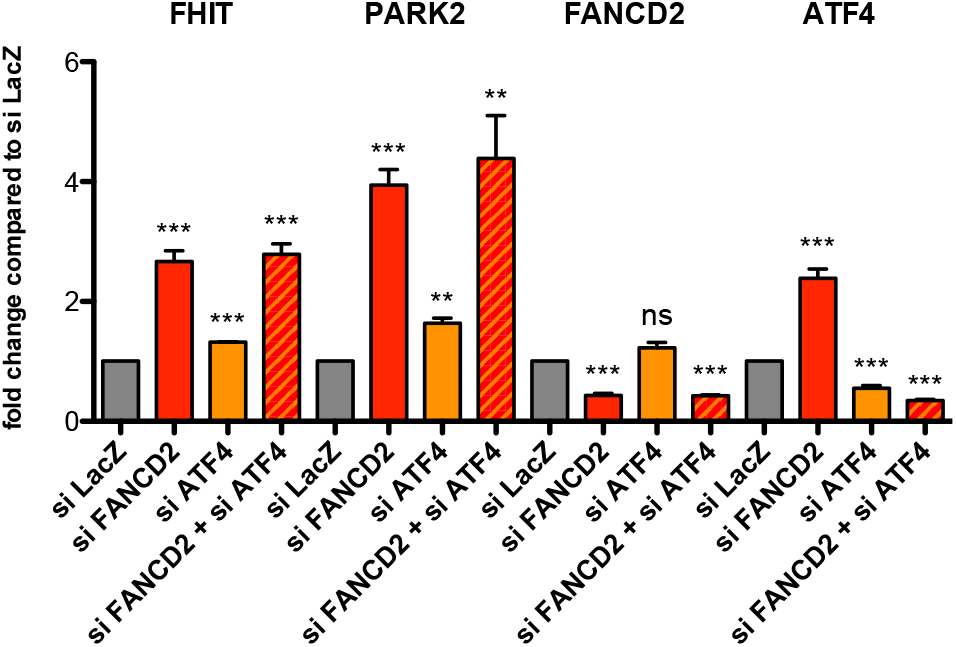
Analysis of CFS gene expression measured by RT-qPCR after control (siLacZ), FANCD2, ATF4, or FANCD2 and ATF4 siRNA transfection. The relative mRNA levels of *FANCD2* and *ATF4* were verified in the same experiments.

**Supplementary Table S1:**
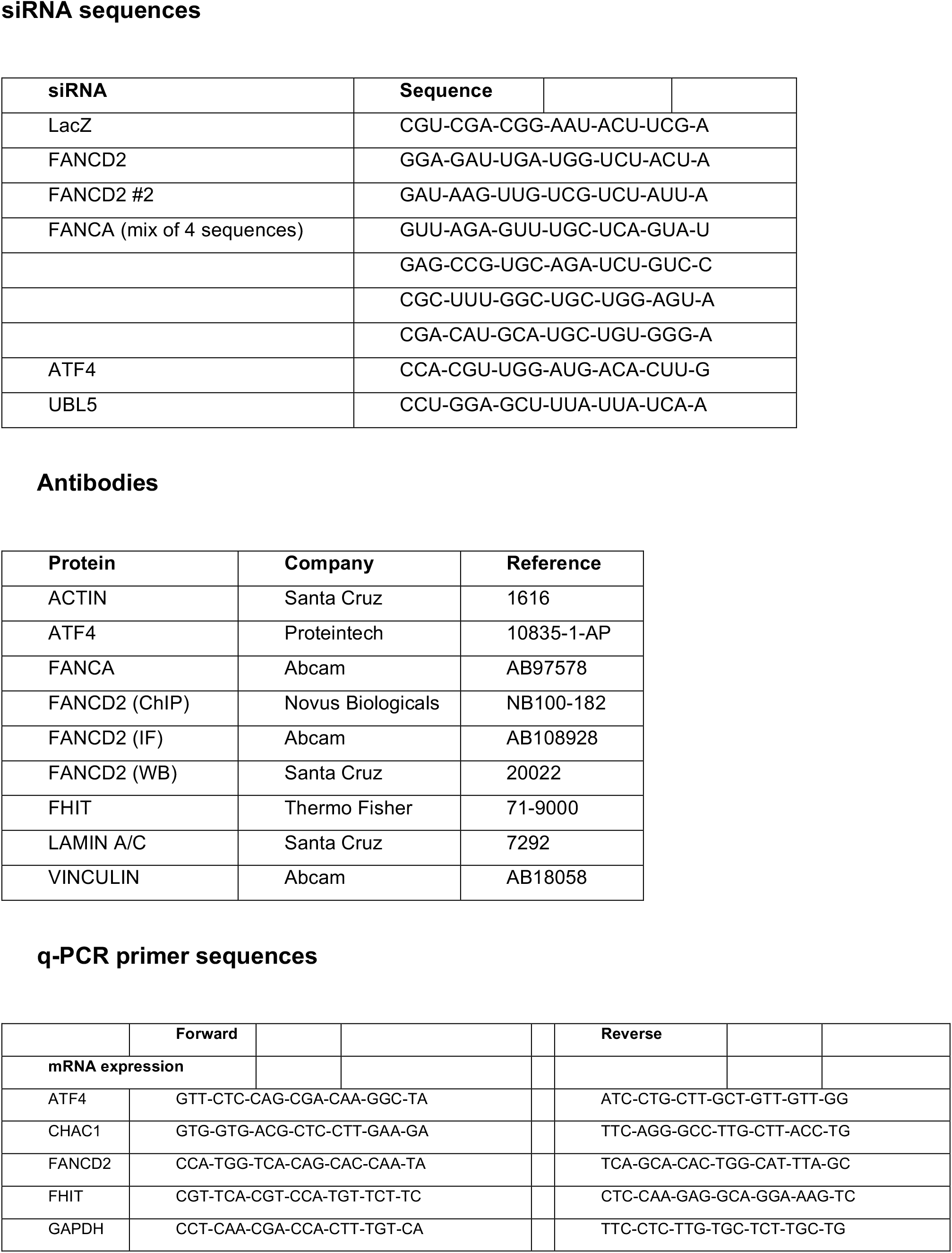
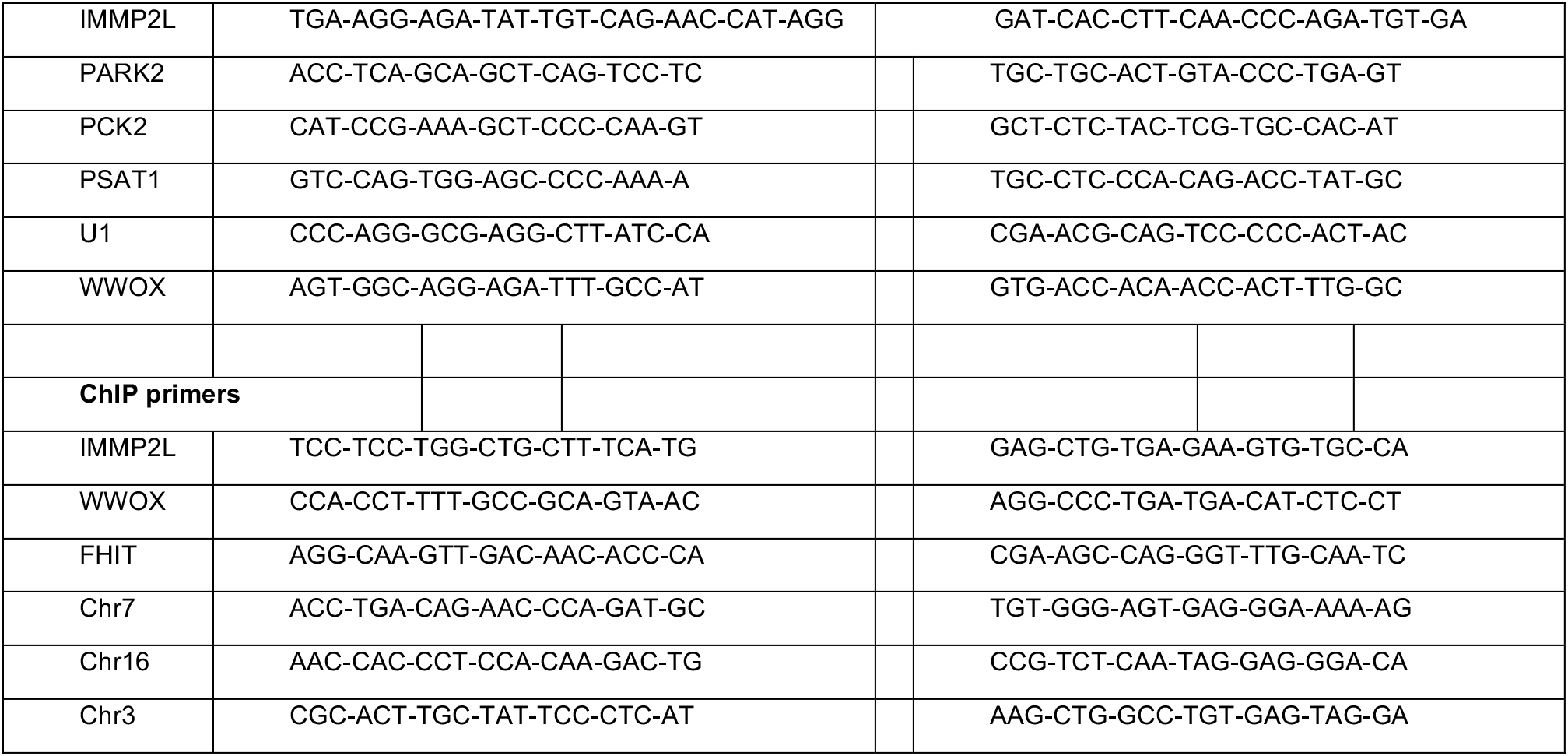
siRNA sequences

